# Improving estimation of species distribution from citizen-science records using data-integration models

**DOI:** 10.1101/2021.04.09.439158

**Authors:** Viviane Zulian, David A. W. Miller, Gonçalo Ferraz

**Affiliations:** Programa de Pós-Graduação em Ecologia, Instituto de Biociências, Universidade Federal do Rio Grande do Sul, CP 15007, Porto Alegre RS 91501-970, Brazil; Department of Ecosystem Science and Management, Pennsylvania State University, 411 Forest Resources Building, University Park, PA 16802, United States

**Keywords:** citizen science, data-integration models, Vinaceous-breasted Parrot, species distribution models, endangered species

## Abstract

Mapping species distributions is a crucial but challenging requirement of wildlife management. The frequent need to sample vast expanses of potential habitat increases the cost of planned surveys and rewards accumulation of opportunistic observations. In this paper, we integrate planned survey data from roost counts with opportunistic samples from eBird, WikiAves and Xeno-canto citizen-science platforms to map the geographic range of the endangered Vinaceous-breasted Parrot. We demonstrate the estimation and mapping of species occurrence based on data integration while accounting for specifics of each data set, including observation technique and uncertainty about the observations. Our analysis illustrates 1) the incorporation of sampling effort, spatial autocorrelation, and site covariates in a joint-likelihood, hierarchical, data-integration model; 2) the evaluation of the contribution of each data set, as well as the contribution of effort covariates, spatial autocorrelation, and site covariates to the predictive ability of fitted models using a cross-validation approach; and 3) how spatial representation of the latent occupancy state (i.e. realized occupancy) helps identify areas with high uncertainty that should be prioritized in future field work. Our results reveal a Vinaceous-breasted Parrot geographic range of 434,670 km^2^, which is three times larger than the ‘Extant’ area previously reported in the IUCN Red List. The exclusion of one data set at a time from the analyses always resulted in worse predictions by the models of truncated data than by the full model, which included all data sets. Likewise, exclusion of spatial autocorrelation, site covariates, or sampling effort resulted in worse predictions. The integration of different data sets into one joint-likelihood model produced a more reliable representation of the species range than any individual data set taken on its own improving the use of citizen science data in combination with planned survey results.

## 1. Introduction

Wildlife management depends on knowledge about species’ geographic ranges, which is also a key element of threat assessment criteria used by the International Union for Conservation of Nature (IUCN, Mace et al. 2008). Despite their unequivocal relevance, accurate range maps are scarce (Jetz et al. 2012). Efforts to improve knowledge about species ranges are hindered by the extent of necessary field sampling and by the scarcity of funding for long-term monitoring. The sampling challenge is heightened by the inevitable tradeoff between data quantity and quality. Planned surveys where observers repeatedly visit a predetermined set of locations using standardized sampling protocols and note the presence or absence of target species provide high-quality information, but they are few and far between. Large and long running planned surveys such as the North American Breeding Bird Survey (BBS; Hudson et al. 2017) or the Pan-European Common Bird Monitoring scheme (PECBM; Gregory et al. 2005) are exceptions to a global pattern of ‘opportunistic’ collection of mostly presence-only data, that records where a species is detected but not where it is searched for and not found.

Technological advances produced many collaborative initiatives where volunteers share wildlife sightings from opportunistic samples in easily accessible online platforms. These initiatives fall under the broad umbrella of citizen-science (Wiggins and Crowston 2011, Tulloch 2013, Heigl et al. 2019). Due to the popularity of birdwatching, citizen-science platforms now hold an extraordinary amount of spatially indexed bird detections. Outstanding examples include the global eBird (Sullivan et al. 2009) and Xeno-canto (Xeno-canto 2019) platforms, as well as the Brazilian WikiAves (WikiAves 2019). These platforms hold data for thousands of bird species, with increasing spatial coverage. These huge datasets have the potential to fill gaps in our knowledge of species’ distributions (Sullivan et al. 2017, Altwegg and Nichols 2018, La Sorte and Somveille 2020). There are, however, wide variations in sampling technique, expertise, and effort among observers, as well as differences in data structures and spatial coverage among citizen-science platforms, which put a premium on the ability to integrate data from different sources. This has spurred progress in the construction of statistical species distribution models that integrate multiple data streams in mapping the probability of species presence over a region of interest (Miller et al. 2019, Fletcher et al. 2019, Isaac et al. 2020).

Initial work on data integration methods focused on presence-only data analyses to discern between areas where a focal species does not occur from areas without sampling where it does occur. Seminal papers by Dorazio (2014), Fithian et al. (2015), and Giraud et al. (2016) integrated presence-only data from opportunistic samples with presence-absence data from planned surveys in a spatial point-process, joint-likelihood framework. The resulting data integration models use the sampling information in presence-absence data to improve inference from the usually larger, presence-only datasets with limited sampling information. This approach has been extended to account for local habitat heterogeneity (Coron et al. 2018) and data patchiness (Peel et al. 2019). In one wide-ranging study, Pacifici et al. (2017) showed how data integration can include site-covariates of species presence, account for spatial autocorrelation, address false positive detections, combine counts with presence-absence data, and weigh datasets differently based on their quality. Simmonds et al. (2020) recently explored the limits of data integration, asking when more data is not necessarily better. These efforts demonstrated how data-integration can not only account for limitations of presence-only data, but also flexibly and robustly harmonize a wide-range of data types (Miller et al. 2019; Isaac et al. 2020).

Development of data integration models emphasized how explicit sampling information from planned-survey, presence-absence data can complement more widely available, opportunistic, presence-only data. Such emphasis may have concealed the extraordinary amount of sampling information in citizen-science datasets themselves. In citizen-science platforms, the set of data points indicating detection of one focal species does not explicitly convey the effort that went into searching for that species. Nonetheless, because platforms gather observations from multiple species, one can find abundant information about sampling effort by looking at where and when non-focal species were detected. Further, citizen-science data frequently include information that can be used to predict sampling effort, such as number of observers, time and distance traveled during sampling, number of detections of all species, or number of species detected. Here, we build on previous work by Fithian et al. (2015), Pacifici et al. (2017), Stauffer et al. (2018), and Miller et al. (2019), to develop a static, integrated occupancy model of species distribution. Our approach assembles detection non-detection histories for each sampling unit and accounts for imperfect detection via the estimation of sampling effort in relation to covariates measured within each citizen-science platform. To assess the extent to which our accounting of sampling effort improves distribution models we employ a cross-validation approach that measures the ability of different models to predict randomly excluded data points. Such assessment of model fit also reveals the extent to which data integration, spatial autocorrelation and site covariates contribute to the modeling task.

Accurate range maps are especially needed for threatened or endangered species in regions that lack planned wildlife surveys, as is often the case in the tropics. The Vinaceous-breasted Parrot (VBP, *Amazona vinacea*) is an endangered species, endemic to the tropical South American Atlantic Forest (BirdLife International 2017). Showing substantial uncertainty about the species’ geographic range, the IUCN reports a ‘possibly extant’ VBP area that is almost three times as large as the ‘extant’ area (Fig. 1A, BirdLife International and Handbook of the Birds of the World 2016). In a recent study of VBP abundance (Zulian et al. 2020) show how ∼75% of known communal roost sites are outside the IUCN ‘extant’ area, suggesting current range estimates are inadequate for planning purposes. This motivated us to ask how existing VBP data sources could be combined to generate a better estimate of the species’ range and identify where the greatest uncertainty in the current distribution exist. We set out to characterize the spatial extent of the current distribution, estimating the local probability of the species’ presence (Kéry 2011) and quantifying the uncertainty about these probability estimates (Rocchini et al. 2011).

**Figure 1.**
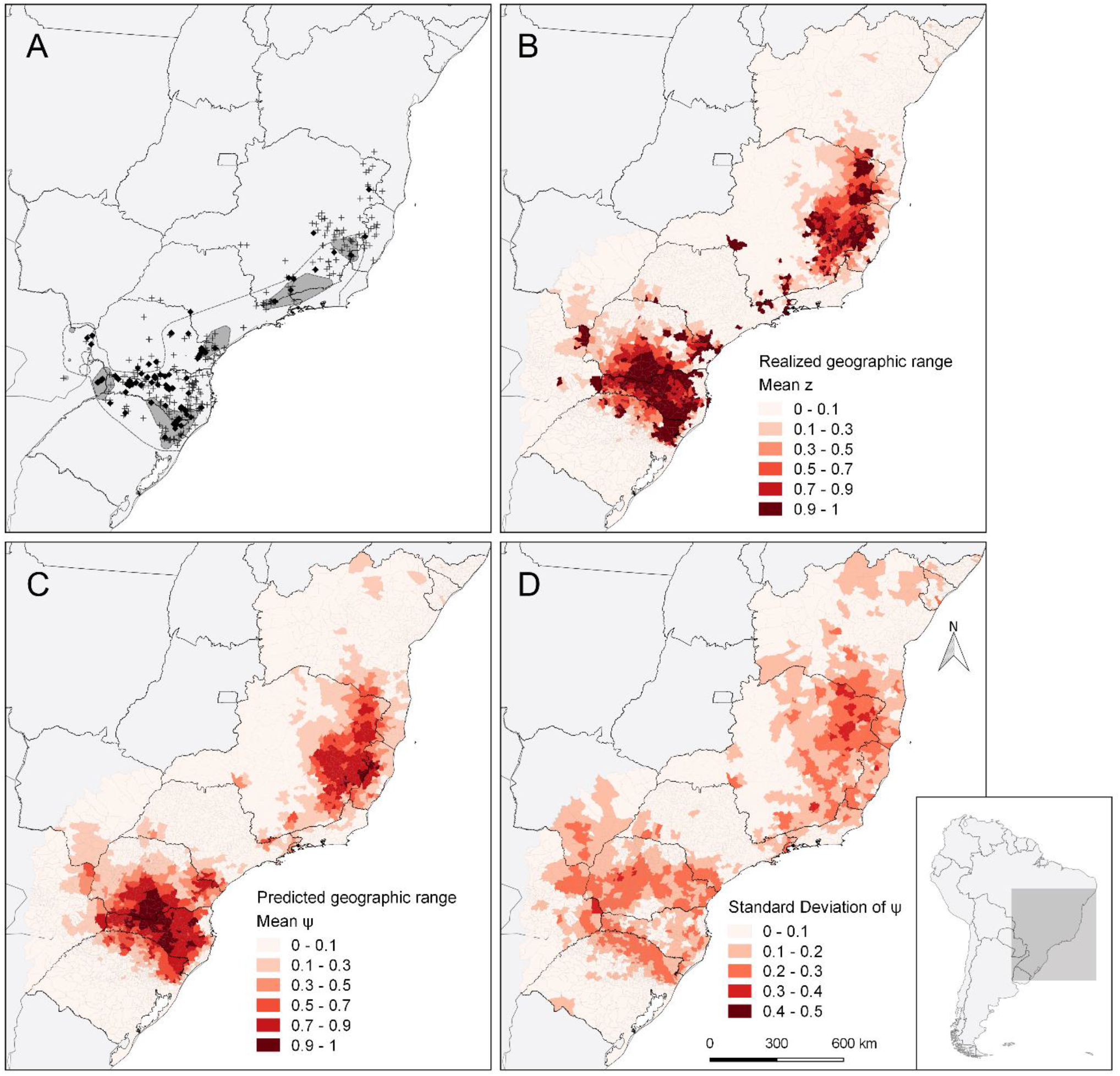
Vinaceous-breasted Parrot observations, geographic distribution, and uncertainty about the distribution. Panel A maps Vinaceous-breasted Parrot detections analyzed in this study with black diamonds indicating the location of roost counts and crosses the location of citizen-science (eBird, WikiAves and Xeno-canto) records. Gray polygons represent the IUCN ‘Extant’ range and dashed lines delimit the IUCN ‘Possibly Extant’ range of the Vinaceous-breasted Parrot. Panel B represents realized occupancy (mean *z*). Panels C and D show, respectively, the predicted occupancy (*ψ*) and the standard deviation of its posterior distribution. Estimates in panels B, C and D are based on the Full Model fit to all (roost counts, eBird, WikiAves and Xeno-canto) datasets. Spatial units correspond to municipalities, with darker tones of red representing higher occupancy (B, C) and higher standard deviation (D).

We aim here to 1) demonstrate how data integration models can be harnessed to address the nuances of data collection across multiple data sets by accounting for variation in sampling effort and detection probability between and within datasets; 2) develop an approach to assess the predictive value of including or excluding different data streams in a single integrated model; and 3) assess how modeling decisions affect the predictive power of our models, with particular attention to the choice of occupancy and detection covariates, whether and how to account for spatial autocorrelation, and how effort and detection are related. We integrate planned-survey data collected by research teams (Zulian et al. 2020) with citizen-science data from the eBird (eBird 2019), WikiAves (WikiAves 2019) and Xeno-canto (Xeno-canto 2019) platforms to model the VBP geographic range in a eleven-year period.

## 2. Methods

### 2.1 Study area

Our study area comprises 2,449,757 km^2^ divided into 3,701 municipalities from Argentina, Brazil, and Paraguay (Fig. 1A). This area includes the entire IUCN-delimited VBP ‘Possibly Resident’ range (BirdLife International and Handbook of the Birds of the World 2016), and is bounded by the limits of the Atlantic Forest biome (Olson et al. 2001). Considering the absence of VBP records north of the Brazilian state of Bahia (BirdLife International 2017), we set the northern limit of our study area along the northern borders of that state and the adjacent state of Alagoas.

### 2.2 Data collection

We obtained VBP detection-nondetection data for all 3,701 municipalities collected between January 1^st^ 2008 and December 31^st^ 2018. The municipality is our spatial sampling unit and we define occupancy as the presence of VBPs in a municipality during the eleven-year study period. Our data come from four sources: roost counts, WikiAves, eBird and Xeno-canto. Roost counts were performed by researchers (Zulian et al. 2020), while WikiAves, eBird, and Xeno-canto data were uploaded to citizen-science platforms by volunteer observers.

Roost count were performed between 2014 and 2018 by 26 teams in 74 municipalities of Brazil, Argentina, and Paraguay, following methodological guidelines described by Zulian et al. (2020). Counts took place between April and June of each year on sites known by researchers to have VBP roosts. Roost count data was converted into detection/non-detection histories with each count from the same municipality considered a replicate sample. Counts with at least one parrot received a ‘1’ (detection) and counts with no parrots received a ‘0’ (nondetection) in the binary history. Parrots are observed in relatively narrow time windows near down and dusk, but early arrivals or a late departures from the roost influence the observations, so we measured the count’s duration in minutes (Time Observing = *TObs*) as an effort covariate.

We obtained eBird data from birding checklists with observations in our study area and uploaded to the platform throughout the study period. All checklists which did not identify a municipality or that potentially spanned more than one municipality due to long distance (>12 km) or long time (>360 minutes) traveled were excluded. Checklists from the same municipality were treated as replicate samples. The checklist structure made it easy to convert eBird data into presence-absence format, and we accordingly built eBird detection/non-detection histories that register the detection (1) or non-detection (0) of the VBP for each list of each municipality. eBird effort covariates were the number of species recorded in a list (*SSee*), minutes spent observing (*TObs*) and kilometers traveled (*RLen*).

WikiAves, receives observer input in the form of individual photos or audio recordings of an identified species and has expert moderators checking uploaded content to avoid misidentification. Record location is registered at the municipality level along with information about authorship and comments. We obtained the total number of records uploaded to each municipality of our study area and period. Our analysis of WikiAves data is based on the set of all records from each municipality and has no replicate samples at the level of the municipality. Thus, there is only one vector of WikiAves detection/non-detection data, with length equal to the number of municipalities and values of ‘1’ or ‘0’, respectively, for those municipalities that did or did not have at least one VBP photo or audio recording. Effort covariates were the number of photos (*NPho*) and audio recordings (*NAud*) submitted to WikiAves per municipality.

Xeno-canto, hosts only audio recordings of bird sounds (Xeno-canto 2019). Unlike WikiAves, Xeno-canto does not have its content checked by moderators, but we did confirm identification of all the VBP records in the Xeno-canto database. We used the R package *warbleR* (Araya-Salas and Smith-Vidaurre 2017) to download the list of all Xeno-canto records from our study area and period. Our Xeno-canto unit data is the set of all audio recordings from one municipality, without replication. We organized these data in the same vector format as WikiAves’ and used the number of recordings (*NAud*) uploaded in each municipality as a covariate of sampling effort.

### 2.3 Data analysis

We summarized each of our four data sources in a matrix or a vector of detection-nondetection information per municipality, depending, respectively, on whether they had multiple (roost counts, eBird) or a single (WikiAves, Xeno-canto) observation per municipality. Effort covariates matrices (or vectors) took the corresponding data source shape. In our models, the true occupancy state of each municipality (or site) *i* is denoted as *z*_*i*_, which takes the value 1 when site *i* was occupied and 0 when not. The state of this latent (partially observed) variable follows a Bernoulli distribution with mean *ψ*_*i*_:

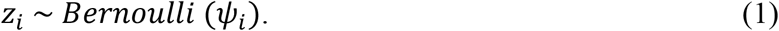

We allowed the probability *ψ*_*i*_ that site *i* is occupied by VBPs to vary with respect to three site environment covariates, according to a generalized linear model with a logit link function. Since VBPs are endemic to the Atlantic Forest and appear to be associated with both altitude (BirdLife International 2017) and Araucaria forest cover (Tella et al. 2016, BirdLife International 2017, Collar et al. 2017, Cockle et al. 2019), we included Atlantic forest cover (*AtF*_*i*_), Araucaria forest cover (*ArF*_*i*_) and average altitude (*Alt*_*i*_) as covariates of municipality *i* occupancy. Forest cover values are from Ribeiro et al. (*in prep*.) as proportions of the municipality area. Average municipality altitude *x*, in meters, is from DIVA-GIS (2018), log-transformed as *log*(*x* + 1). Our linear model of occupancy also included a spatial random effect to account for unexplained spatial autocorrelated variation (*δ*_*i*_):

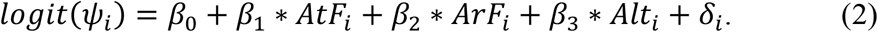

This effect follows a conditional auto-regressive (CAR) distribution as applied by Pacifici et al. (2017) in the context of integrated species distribution models. To avoid confounding effects of municipality size variability and to gain replication among spatial units in the CAR analysis, we represented space by an hexagonal lattice overlaid on the study area, with municipalities assigned to the lattice cell that matches their centroid. Cells measured 0.5° latitude across; all the first-order neighbors of each cell were given a weight of 1 when fitting the CAR model.

We fit a joint-likelihood data-integration model with a single shared occupancy process: for all four data types, VBP detection in sample *j* and site *i* is conditional on the species being present at the site (*z*_*i*_ = 1). Departing slightly from the standard accounting of effort based on the number of replicate samples (MacKenzie et al. 2002), we express the conditional probability 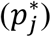 of detecting the species as a function of an estimated amount of sampling effort (*E*_*j*_) for visit *j* (Stauffer et al. 2018, Miller et al. 2019):

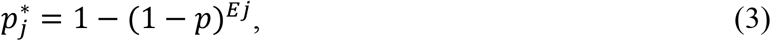

where *p* is the probability of detection per unit effort. Because we are using indirect, and sometimes several metrics of effort for each data source (our effort covariates), we estimate parameter *E*_*j*_ for each sample *j* as a linear function of the covariates. Thus, for each data source (RC=roost counts, EB=eBird, WA=WikiAves and XC=Xeno-canto) we have:

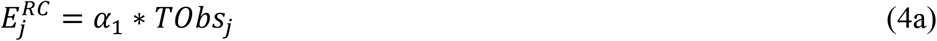

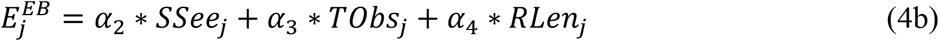

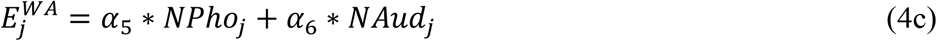

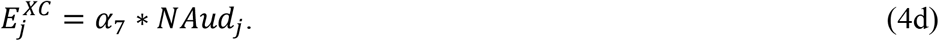

Equations 4a-d have no intercept, so that effort is 0 when all effort covariates are 0. In addition, we fix *p* at a value of 0.5, so that the *α*_1−7_ coefficients express the relationship between covariates and the effort necessary to reach a detection probability of 0.5. Without fixing *p*, equation 3 becomes over-parameterized. Coefficients *α*_1−7_ of the effort functions also show the relative contribution of each effort metric to the total estimated effort per dataset (see code in Supplemental Material S1). Finally, our detection/non-detection histories *Y*_*ij*_ in each dataset follow the Bernoulli distribution:

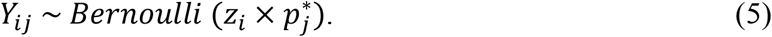

We first fitted a Full Model accounting for the effects of all effort metrics, all site covariates, and spatial autocorrelation. Subsequently, we evaluated the impact of different modeling decisions on predicted accuracy by fitting seven additional ‘incomplete’ models listed in Table 2. We fitted all the models using a Bayesian estimator coded in the BUGS language and run on WinBugs software (Lunn et al. 2000), which includes predefined model structures for CAR random effects. Inference was based on draws from the posterior distribution of model parameters using an MCMC algorithm with three chains, 200,000 iterations, and a burn-in phase of 100,000. We considered parameters with an R-hat lower than 1.1 to have converged and used results to draw parameter posterior distributions.

We accessed model fit by excluding all the detection non-detection data from a randomly selected set of 650 municipalities (20% of the total), fitting the models to the training dataset (i.e. truncated data) and then predicting the validation dataset (excluded data) based on the estimated parameters. In this cross-validation approach, our predicted accuracy measures a model’s ability to predict excluded data as expressed by the likelihood-based Deviance:

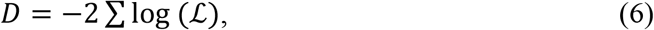

where the likelihood *ℒ* equals *ŷ*^*y*^ ∗ (1 − *ŷ*)^1−*y*^ for each site and visit in the validation dataset (Hooten and Hobbs 2015). We use *y* and *ŷ* to represent, respectively, the observed, binary data and the predicted probability of detecting VBPs for each site and visit based on estimates from the training data. The lowest deviance values indicate the best fit. We examined overall model deviance, summed across data sources, as well as individual deviance values for each data source to look at source-specific predictive performance. Comparisons among values revealed the impact of site covariates, detection covariates, or the CAR component on the predictive performance of our models.

To determine whether each of the individual data sets improved the predictive ability of our model, we fit the model to four truncated data sets, including all covariates and the CAR random effect, but excluding one data source at a time. Such rotating exclusion made it possible to examine whether the addition of a data source to the mix improves the model’s ability to predict the validation set from other sources. Specifically, we asked whether predictions of validation data from a training data source, were more or less accurate when each of the other data sources were excluded. For example, if eBird does contribute to improving the overall model, then including eBird data should lead to better predictions of Xeno-canto, WikiAves, and roost count data. This is a measure of overall prediction consistency among data sources.

Finally, we represent the VBP geographic range using two estimates of site-occupancy. The first, ‘realized’ occupancy, is conditional on the observations; it equals 1 in all municipalities where VBP was seen at least once, and is the expected value of the latent occupancy state (*z*_*i*_) where it was not seen. As effort increases and VBP are not observed, *z* converges toward 0, and so does realized occupancy. This metric provides a measure of local uncertainty about species presence given all available data and, unlike typical predictions by distribution models, accurately expresses local certainty of occurrence by adjusting predictions to actual observation. The second estimate, ‘predicted’ occupancy, is not conditioned on the actual data for a municipality. It offers estimates of *ψ*_*i*_, which express occupancy probability for a statistical population of municipalities with the same site covariates and neighborhood of municipality *i* (Fig. 1C). Predicted occupancies are typically visualized in distribution models, expressing how estimated environmental relationships affect the local probability of occurrence across a species range.

## 3. Results

We draw on 1,007 VBP detections from 47,240 samples in four datasets collected across the 3,402 municipalities within our study area. While the roost count data contains 40% of all detections, roost counts covered only 2.2% of the municipalities in our study area. The 596 detections returned by citizen-science platforms, on the other hand, come from 3,401 municipalities (Table 1). WikiAves had the widest coverage, with data for 3,190 municipalities, and VBP detections for 191 of them. eBird had smaller coverage than WikiAves but had the largest number of samples of all sources, with 388 VBP detections from 71 municipalities. The number of eBird samples varied substantially across municipalities, ranging from 1 to 3,244 (São Paulo, SP, Brazil) with a mean of 42. Xeno-canto had the smallest coverage and number of detections. The highest detection rate—given by the ratio of *n*_det_ to Sample size, in Table 1— appears in the roost count dataset (88%), as expected, because the sampling focused in locations where VBPs were known to occur. The dataset with the highest number of samples (eBird) had the lowest detection rate, of 1%.

**Table 1.**
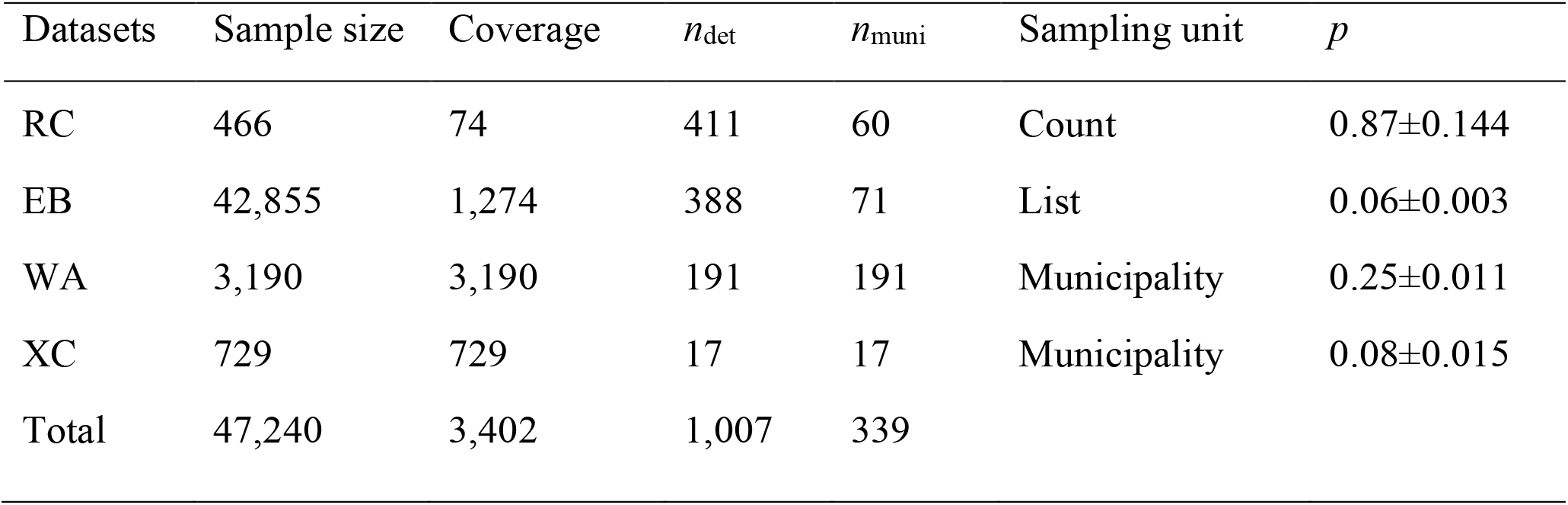
Sample size, spatial coverage, and number of Vinaceous-breasted Parrot detections from roost counts (RC), eBird (EB), WikiAves (WA) and Xeno-canto (XC). Sample size is number of samples, following each database’s sampling unit definition. Spatial coverage is the number of municipalities sampled, with total smaller than the sum across databases because some municipalities are included in more than one database. Labels *n*_det_ and *n*_muni_ show, respectively, the number of parrot detections and the number of municipalities with at least one detection. The sampling unit is the data category considered as a replicate; *p* is estimated detection probability per sampling unit at average effort for each dataset, under the Full Model.

| Datasets | Sample size | Coverage | $n_{\text{det}}$ | $n_{\text{muni}}$ | Sampling unit | $p$ |
| --- | --- | --- | --- | --- | --- | --- |
| RC | 466 | 74 | 411 | 60 | Count | 0.87±0.144 |
| EB | 42,855 | 1,274 | 388 | 71 | List | 0.06±0.003 |
| WA | 3,190 | 3,190 | 191 | 191 | Municipality | 0.25±0.011 |
| XC | 729 | 729 | 17 | 17 | Municipality | 0.08±0.015 |
| Total | 47,240 | 3,402 | 1,007 | 339 |  |  |

Tabel 2 shows model predictive ability based on cross-validation. The Full Model had the best predictive ability. The exclusion of detection covariates (Model 3) had the greatest negative impact on predictive ability, with estimated deviance being 2.15 times higher for this model than for the Full Model (Table 2). Despite producing the highest Total Deviance with combined data sets, the exclusion of detection covariates in Model 3 improved the fit to Xeno-canto data. Model 2, which does not account for spatial autocorrelation, had the second worse predictive ability, followed by Model 4, which did not include covariates for occupancy. Despite having higher Total Deviance than the Full Model, Model 4 showed the best fit to roost count data, of all models (Table 2). Our results reveal that effort-based modeling of detection, inclusion of spatial auto-correlation in occupancy, and consideration of environmental covariates improved the predictive ability of our species distribution models.

**Table 2.** Deviance for each site-occupancy model in this study. Model 1, designated as ‘Full Model’, includes detection as well as occupancy covariates and was fitted to data from all datasets: roost counts (RC), eBird (EB), WikiAves (WA), and Xeno-canto (XC). Model 2 equals model 1 without spatial autocorrelation. Models 3 and 4 are variants of model 1 without, respectively, detection and occupancy covariates. Models 5-8 differ from the Full Model by the exclusion of one data set each, as shown. Since models 5-8 do not use the same data, their Total Deviance is not comparable and is omitted from the Table. Bold font highlights the model with the best fit by Total Deviance.

| Models | Deviance in each data set |  |  |  |  |
| --- | --- | --- | --- | --- | --- |
|  | Total Deviance | RC | EB | WA | XC |
| **1. Full Model** | **440.85** | **28.84** | **281.19** | **103.35** | **27.46** |
| 2. No CAR | 581.32 | 50.97 | 362.58 | 139.56 | 28.20 |
| 3. No detection covs. | 952.84 | 57.21 | 735.61 | 133.60 | 26.41 |
| 4. No occupancy covs. | 477.06 | 26.04 | 315.34 | 107.79 | 27.87 |
| 5. All data but RC | — | — | 301.88 | 108.78 | 27.56 |
| 6. All data but EB | — | 28.53 | — | 110.26 | 25.55 |
| 7. All data but WA | — | 35.27 | 326.01 | — | 28.61 |
| 8. All data but XC | — | 34.04 | 309.29 | 116.18 | — |

Models 5-8 assess whether individual data sets improve overall predictive ability. We compare dataset-specific deviances from the validation data for each of the four models to that of the full model (where no data was excluded). Including the four data sets in the analysis (i.e., using the Full Model) improved fit in all but two cases. Dataset-specific deviances of Models 5, 7, and 8 were all higher—indicating lower prediction power— than those of the Full Model (Table 2). Removal of eBird data (Model 6) slightly improved the prediction of Xeno-canto data, but clearly worsened the fit to WikiAves data, leaving that of the roost count data virtually unchanged.

The sum of municipality areas weighted by the Full Model realized-occupancy estimates returned a realized VBP range of 434,670 km^2^, which is three times larger than the IUCN Red List ‘Extant’ area (BirdLife International and Handbook of the Birds of the World 2016). Both the realized and the predicted ranges appear split in two large patches (Fig. 1 B and C). The southern patch covers parts of Argentina, Paraguay, and the Brazilian states of Rio Grande do Sul, Santa Catarina and Paraná; the northern patch overlaps the Brazilian states of Minas Gerais, Espírito Santo and Bahia. The realized range also includes small areas between the two large patches, mainly in the Campos do Jordão region, near the border between São Paulo and Minas Gerais (Fig. 1B). Uncertainty about the VBP range is greatest around high-occupancy patch edges, as shown by intermediate values of realized occupancy (Fig. 1B) and high standard deviation of predicted occupancy (Fig. 1D). As expected, municipalities with the most extreme occupancy values (close to 0 or 1) returned the lowest standard deviation values (Fig. 1D).

Araucaria and Atlantic forest cover had strong (and positive) effects on occupancy probability (Table 3). Altitude had a weaker, positive, but more precise effect on *ψ*, when compared with the two forest covariates. The different effort covariates on the bottom part of Table 3 had varying, though always positive, effects on detection probability. Among these, time spent observing showed the highest effect, both as α_1_, which measures the duration of a roost count, and as α_3_, the time spent collecting an eBird list. The number of audio recordings uploaded in WikiAves was a stronger predictor of survey effort (α_6_), and thus overall detection probability for a municipality, than the number of photos (with effect α_5_). Detection probability at the average effort was highest for roost counts, carried out at known parrot roosts, with a posterior mean and standard deviation of 0.87 ± 0.144. Mean detection probability among citizen-science datasets was highest for WikiAves (0.25 ± 0.011), and lowest for eBird (0.06 ± 0.003) and Xeno-canto (0.08 ± 0.015; Table 1).

**Table 3.** Estimated mean, standard deviation (SD) and 95% credible intervals (CI) for the posterior distribution of Full Model coefficients. Occupancy function coefficients (*β*_1_ to σ) specify the biological process, while detection coefficients (α_1_–α_7_) specify the sampling process. The covariates corresponding to each coefficient appear in parentheses in front of its name; σ measures the magnitude of spatial autocorrelation in site occupancy. Coefficients α_1_, α_2_–α_4_, α_5_– α_6_ and α_7_ correspond, respectively, to metrics of effort per municipality in roost counts (RC), eBird (EB), WikiAves (WA), and Xeno-canto (XC) databases. Each metric is indicated in parentheses in front of the coefficient name.

| *Biological Process* |  |  |
| --- | --- | --- |
| Parameter | Mean ± SD | 95% CI |
| $\beta_1$ (Altantic forest cover) | 2.110 ± 0.8684 | 0.379 – 3.792 |
| $\beta_2$ (Araucaria forest cover) | 2.133 ± 0.9806 | 0.296 – 4.104 |
| $\beta_3$ (Altitude) | 0.852 ± 0.1205 | 0.579 – 1.055 |
| *Sampling Process* |  |  |
| Parameter | Mean ± SD | 95% CI |
| $\alpha_1$ (RC: Time observing) | 1.814 ± 0.1110 | 1.613 – 2.043 |
| $\alpha_2$ (EB: Species seen) | 0.002 ± 0.0002 | 0.001 – 0.002 |
| $\alpha_3$ (EB: Time observing) | 0.008 ± 0.0027 | 0.003 – 0.013 |
| $\alpha_4$ (EB: Route length) | 0.005 ± 0.0019 | 0.002 – 0.009 |
| $\alpha_5$ (WA: Photos) | 0.001 ± 0.0002 | 0.001 – 0.002 |
| $\alpha_6$ (WA: Audio recordings) | 0.006 ± 0.0021 | 0.003 – 0.011 |
| $\alpha_7$ (XC: Audio recordings) | 0.007 ± 0.0017 | 0.004 – 0.011 |

## 4. Discussion

The Vinaceous-breasted Parrot geographic range covers approximately 434 thousand square kilometers subdivided into two large patches, one centered in the southern Brazilian state of Santa Catarina and another to the north, centered in eastern Minas Gerais state, also in Brazil. A third, much smaller area of occupancy comprises a group of relatively high-altitude municipalities near Campos do Jordão, in São Paulo and Minas Gerais states, approximately 100 kilometers west of the Rio de Janeiro border. Our two-patch range contrasts with the five patches represented in the IUCN ‘resident’ range. The ‘possibly resident’ IUCN range, which encloses all of the ‘resident’ patches, conveys uncertainty about the subdivision in five areas (BirdLife International and Handbook of the Birds of the World 2016). Our study provides evidence for redrawing the VBP range while quantifying uncertainty associated with the new map. We look forward to seeing population genetic studies that elucidate the extent of reproductive isolation between the two large patches, as well as between the small Campos do Jordão area and the northern, Minas Gerais patch. A comparison between realized and predicted ranges shows that some municipalities with high mean *z* have relatively low predicted occupancy probability (*ψ*). Some VBP observations in these municipalities may correspond to animals released or escaped from captivity, but these locations deserve further investigation, particularly those in south-west Minas Gerais and south-west São Paulo, to exclude the possibility of there being unknown isolated populations. Intermediate values of realized occupancy and high standard deviation of the posterior distribution of predicted occupancy reveal areas with high uncertainty about VBP presence, which, like the isolated high-*z* municipalities, ought to be targeted by future field searches. Three regions stand out for high uncertainty about VBP presence: northeastern Minas Gerais, central Paraná, and northern Rio Grande do Sul, in Brazil, together with a few municipalities in eastern Paraguay. These are the regions that could contribute most to further improvement of knowledge about the VBP geographic range.

Our estimated VBP range exceeds the area of past Araucaria forest mapped by Hueck (1966) and includes vast areas of the Atlantic forest biome that have been cleared. Nonetheless, both vegetation site-covariates—Araucaria and Atlantic forest cover—had strong positive effects on site-occupancy probability. The Paraná Pine plays an important role in the VBP natural history, at least in part of its range, offering roost sites (Prestes et al. 2014), nesting cavities (Cockle et al. 2007), and nutrition during the coldest months of the year (Prestes et al. 2014, Kilpp et al. 2015, Tella et al. 2016, Collar et al. 2017). Nevertheless, since Araucaria forests only extend as far north as the Campos do Jordão region, parrots from the northern patch must rely on other plant species to obtain whatever resources their southern counterparts get from the Paraná Pine. Living at a lower latitude, they may also escape the harshness of cold winter weeks, when Araucaria seeds are a unique source of energy for several species of the southern fauna (Dénes et al. 2018). Indeed, Carrara et al. (2008) registered foraging and roosting in different trees between northern and southern locations. Likewise, Cockle et al. (2007), as well as Prestes et al. (2014), document foraging and cavity nesting in non-Araucaria Atlantic Forest trees of the southern part of the range. The effect of altitude on site-occupancy was smaller and more uncertain than the effects of forest cover, but still indisputably positive. Thus, environmental consequences of altitude are not limiting the VBP distribution.

The increasing availability of citizen-science datas offers a great opportunity to improve species distribution maps. In our study, eBird, WikiAves, and Xeno-canto jointly produced 1.45 times more VBP detections, from samples that covered 45 times more municipalities, than the researcher-led counts. The integration of four different datasets substantially increased spatial coverage—improving accuracy—and, by enlarging sample size, resulted in improved precision of occupancy estimates. The assessment of predictive accuracy enabled us to measure the contribution of each dataset for the final estimates. Excluding one data set at a time from the analyses always resulted in worse prediction by the truncated analysis than by the joint analysis of all data sets. Exclusion of the WikiAves data had the highest impact on predictive power, increasing the Deviance of the other data sets between 4% and 25%. WikiAves still lacks an automated data download tool, but it is currently the best source of bird species distribution information in the Brazil because it has the best coverage and highest number of records among available citizen-science sources. Xeno-canto has the fewest records and smallest spatial coverage, but it still produced a measurable improvement of predicting power when added to the other data sets. Roost counts and eBird had the least consistent impact on prediction power but still produced an average decrease in deviance across datasets. These two datasets also contributed with sampling replication, essential for the quantification of false negative results. While there are limits to the usefulness of data integration (Simmonds et al. 2020), in our case, integration clearly improved the fit of models, suggesting that different datasets are capturing similar realities of parrot distribution; otherwise, their combination should make it more, not less difficult to predict excluded data. Comparisons across datasets were only possible thanks to a methodology that explicitly accounts for differences in data collection and observation processes.

Exclusion of the occupancy covariates and the spatial autocorrelation component increased total deviance by 8 and 31%, respectively. The strong effect of spatial autocorrelation on deviance signals a spatially structured geographic distribution. Neglecting spatial contagion easily leads to biased parameter estimates, potentially resulting in erroneous distribution maps (Johnson et al. 2013, Guélat and Kéry 2018). We emphasize that even though our sampling units varied substantially in size, spatial effects can be interpreted equally throughout the study area because we estimated spatial autocorrelation (CAR) parameters over an hexagonal grid with fixed-size cells.

The term ‘citizen science’ covers a wide variety of collaborative arrangements that involve people from outside the scientific community in scientific research (Wiggins and Crowston 2011, Tulloch 2013, Heigl et al. 2019). When it comes to collaborative recording of wildlife sightings, however, most citizen-science initiatives compile presence-only information from opportunistic samples. Our analysis employs presence-absence (roost counts, eBird) and presence-only (WikiAves, Xeno-canto) data, as well as a planned survey (roost counts) and opportunistic sampling (WikiAves, eBird, Xeno-canto). Our model does integrate planned-survey with opportunistic sampling data, but we want to emphasize the approach to accounting for spatial bias in citizen-science data: the estimation of effort based on multiple covariates that can be obtained from the citizen-science datasets themselves (Miller et al. 2019). This approach is synthesized in equations 3 and 4a-c, which express detection probability conditional on species presence. Total deviance more than doubled when effort covariates were removed from the analysis, but it increased only up to 7% (for the eBird data) when we removed the planned-survey roost count data. This result is in agreement with the usefulness of integrating citizen-science with planned survey data, and it strengthens our confidence in the contribution of large, multi-species citizen-science datasets for improving knowledge about species distributions.

## Acknowledgements

This paper owes a great deal to Reinaldo Guedes who volunteered his free time over the last twelve years to developing and administrating WikiAves, the most successful citizen-science initiative in Brazil. Roost counts were supported in large part by *Projeto Charão*, in Brazil, *Proyecto Selva Pino Paraná*, in Argentina, and *Guyra Paraguay*, in Paraguay. Last, but not least, we are indebted to the thousands of bird observers who uploaded photos, audio recordings and birding lists to eBird, WikiAves and Xeno-canto.

