## Supplemental Material S1 for "Improving estimation of species distribution from citizen-science records using data-integration models"

**Supplemental Material Appendix A:** R and BUGS code for the Full Model used in estimating the Vinaceous-breasted Parrot geographic range. The code includes the CAR component of the model, accounting for effort, cross-validation and Deviance computation.

```
#Data object specification in R
data <- list(muni1 = datIn$IDENT, muni2 = datOut$IDENT, muni3 = dat2In$IDENT,
            muni4 = dat2Out$IDENT, muni5 = dat3In$IDENT, muni6 =
            dat3Out$IDENT, muni7 = dat4In$IDENT, muni8 = dat4Out$IDENT,
            Y1 = datIn$A_VINACEA, Y3 = dat2In$A_VINACEA, Y5 =
            dat3In$AVINACEA, Y7 = dat4In$A_VINACEA, nMuni = length(muni),
            nObs1 = nrow(datIn), nObs2 = nrow(datOut), nObs3 = nrow(dat2In),
            nObs4 = nrow(dat2Out), nObs5 = nrow(dat3In), nObs6 =
            nrow(dat3Out), nObs7 = nrow(dat4In), nObs8 = nrow(dat4Out),
            TObs = datIn$EFFORT_MIN/60, TObs2 = datOut$EFFORT_MIN/60,
            SSee = dat2In$NSPECIES, SSee2 = dat2Out$NSPECIES, TObs3 =
            dat2In$DURATION.M/60, TObs4 = dat2Out$DURATION.M/60,
            RLen = dat2In$EFFORT.DIS, RLen2 = dat2Out$EFFORT.DIS,
            NPho = dat3In$NPIC, NPho2 = dat3Out$NPIC, NAud = dat3In$NSONG,
            NAud2 = dat3Out$NSONG, NAud3 = dat4In$NSONGS, NAud4 =
            dat4Out$NSONGS, VegCover = VegCover, ArauCover = ArauCover,
            Alt = Altitude, nCell = nrow(hex.centroids), cell.id = cell.id,
            adj = adj, num = num, sumNeigh = sumNeigh)

# Model specification in BUGS language
cat(file = "model.txt","
model {

#CAR prior - spatial random effect
for(j in 1:sumNeigh){weights[j] <- 1}
spacesigma ~ dunif(0,5)
spacetau <- 1/(spacesigma*spacesigma)
delta[1:nCell] ~ car.normal(adj[],weights[],num[],spacetau)

###Data model
for (i in 1:nMuni){   #loop over sites
    mu[i] <- delta[cell.id[i]] + beta[1] + beta[2]*VegCover[i] +
             beta[3]*ArauCover[i] + beta[4]*Alt[i]
    mu.lim[i] <- min(10, max(-10, mu[i]))
    logit(psi[i]) <- mu.lim[i]
    z[i] ~ dbern(psi[i])
}

for (n in 1:nObs1){      #loop over observations – Count Data - Data In
    e1[n] <- alpha[1]*TObs[n]
    P1[n] <- 1-pow((1-0.5), e1[n])
    zP1[n] <- P1[n]*z[muni1[n]]
    Y1[n] ~ dbern(zP1[n])
}

for (o in 1:nObs2){      #loop over observations – Count Data - Data Out
    e2[o] <- alpha[1]*TObs2[o]
    P2[o] <- 1-pow((1-0.5), e2[o]) #effort model
```

```
    zP2[o] <- P2[o]*z[muni2[o]]
    Y2[o] ~ dbern(zP2[o])
}

for (j in 1:nObs3){    #loop over observations - eBird Data
    e3[j] <- alpha[2]*SSee[j] + alpha[3]*TObs3[j] + alpha[4]*RLen[j]
    P3[j] <- 1-pow((1-0.5), e3[j]) #effort model
    zP3[j] <- P3[j]*z[muni3[j]]
    Y3[j] ~ dbern(zP3[j])
}

for (p in 1:nObs4){  #loop over observations - eBird Data - Cross-Validation
    e4[p] <- alpha[2]*SSee2[p] + alpha[3]*TObs4[p] + alpha[4]*RLen2[p]
    P4[p] <- 1-pow((1-0.5), e4[p]) #effort model
    zP4[p] <- P4[p]*z[muni4[p]]
    Y4[p] ~ dbern(zP4[p])
}

for (k in 1:nObs5){  #loop over observations - Wikiaves data
    e5[k] <- alpha[5]*NPho[k] + alpha[6]*NAud[k]
    P5[k] <- 1-pow((1-0.5), e5[k]) #effort model
    zP5[k] <- P5[k]*z[muni5[k]]
    Y5[k] ~ dbern(zP5[k])
}

for (s in 1:nObs6){  #loop over observations - Wikiaves data - Cross-
Validation
    e6[s] <- alpha[5]*NPho2[s] + alpha[6]*NAud2[s]
    P6[s] <- 1-pow((1-0.5), e6[s]) #effort model
    zP6[s] <- P6[s]*z[muni6[s]]
    Y6[s] ~ dbern(zP6[s])
}

for (h in 1:nObs7){  #loop over observations - Xeno-Canto data
    e7[h] <- alpha[7]*NAud3[h]
    P7[h] <- 1-pow((1-0.5), e7[h]) #effort model
    zP7[h] <- P7[h]*z[muni7[h]]
    Y7[h] ~ dbern(zP7[h])
}

for (t in 1:nObs8){  #loop over observations - Xeno-Canto - Cross-Validation
e8[t] <- alpha[7]*NAud4[t]
    P8[t] <- 1-pow((1-0.5), e8[t]) #effort model
    zP8[t] <- P8[t]*z[muni8[t]]
    Y8[t] ~ dbern(zP8[t])
}

#Priors for betas - psi
beta[1] ~ dunif(-10,10)
beta[2] ~ dunif(-10,10)
beta[3] ~ dunif(-10,10)
beta[4] ~ dunif(-10,10)
```

```
#Priors for alphas - effort model
for (b in 1:7){
    alpha[b] ~ dnorm(0,0.0001)I(0,10000)
}

#compute the mean detection probability of each dataset:
muP1 <- mean(P1[])
muP2 <- mean(P2[])
muP3 <- mean(P3[])
muP4 <- mean(P4[])
muP5 <- mean(P5[])
muP6 <- mean(P6[])
muP7 <- mean(P7[])
muP8 <- mean(P8[])

}
")

#Back to R language:

#Specification of Initial Values
inits = function() {list(z = rep(1, data4$nMuni))}
params <- c("beta", "psi", "z", "alpha", "muP1", "muP2", "muP3", "muP4",
           "muP5", "muP6", "muP7", "muP8", "Y2", "Y4", "Y6", "Y8",
           "spacesigma", "delta")

#MCMC settings
nc <- 3;   nb <- 150000;   ni <- 200000;   nt <- 100

out <- bugs(data = data, inits = inits, parameters.to.save = params,
           model.file = "model.txt", n.chains = nc, n.iter = ni,
           n.burnin = nb, n.thin = nthin, debug = TRUE)

#Deviance calculation based on the model output
likhood2 <- (out$mean$Y2^datOut$A_VINACEA)*((1- out$mean$Y2)^
             (1-datOut$A_VINACEA)) #likelihood
DEV2 <- -(2*(sum(log(likhood2))))

likhood4 <- (out$mean$Y4^ dat2Out$A_VINACEA)*((1- out$mean$Y4)^
             (1- dat2Out$A_VINACEA)) #likelihood
DEV4 <- -(2*(sum(log(likhood4))))

likhood6 <- (out$mean$Y6^ dat3Out$AVINACEA)*((1- out$mean$Y6)^
             (1- dat3Out$AVINACEA)) #likelihood
DEV6 <- -(2*(sum(log(likhood6))))

likhood8 <- (out$mean$Y8^ dat4Out$A_VINACEA)*((1- out$mean$Y8)^
             (1- dat4Out$A_VINACEA)) #likelihood
DEV8 <- -(2*(sum(log(likhood8))))

DEVtotal <- DEV2 + DEV4 + DEV6 + DEV8 #total deviance
```
